# Evolutionary convergence of a neural mechanism in the cavefish lateral line system

**DOI:** 10.1101/2022.01.26.477913

**Authors:** Elias T. Lunsford, Alexandra Paz, Alex C. Keene, James C. Liao

## Abstract

Animals can evolve dramatic sensory functions in response to environmental constraints, but little is known about the neural mechanisms underlying these changes. The Mexican tetra, *Astyanax mexicanus*, is a leading model to study genetic, behavioral, and physiological evolution by comparing eyed surface populations and blind cave populations. We compared neurophysiological responses of posterior lateral line afferent neurons and motor neurons across *A. mexicanus* populations to reveal how shifts in sensory function may shape behavioral diversity. These studies indicate differences in intrinsic afferent signaling and gain control across populations. Elevated endogenous afferent activity identified a lower response threshold in the lateral line of blind cavefish relative to surface fish. We next measured the effect of inhibitory corollary discharges from hindbrain efferent neurons onto afferents during locomotion. We discovered that three independently-derived cavefish populations have evolved persistent afferent activity during locomotion, suggesting for the first time that regression of the efferent system can be an evolutionary mechanism for neural adaptation of a vertebrate sensory system.

## Introduction

Our understanding of the sensory systems and behavior of animals is challenging to contextualize within the framework of evolution. Anatomical comparisons between species have allowed us to infer sensory capabilities, but this approach cannot directly reveal neural function. Discovery of neural mechanisms that underlie behavior are often constrained to a limited number of model species (Jourjine & Hoekstra, 2021). Like morphology, neural circuits can adapt to the environment. Of these, many circuits are sensory and regulate essential behaviors such as foraging, navigation, and escapes (Blin et al., 2018; Hoke et al., 2012; Hüppop, 1987; Paz et al., 2020).

The Mexican blind cavefish, *Astyanax mexicanus*, is a powerful model to understand the evolution of physiological and molecular traits that contribute to behaviors such as sleep (Duboué et al., 2011; J. B. Jaggard et al., 2018) and prey capture (Lloyd et al., 2018; Yoshizawa et al., 2014). *Astyanax mexicanus* exists in two morphs; 1) eyed surface-dwelling populations and 2) blind cave populations. There are at least 30 independently evolved cavefish populations in the caves of the Sierra de El Abra region of Northeast Mexico (Herman et al., 2018; McGaugh et al., 2020; RW Mitchell et al., 1997). *Astyanax mexicanus* populations are interfertile, and this attribute has allowed investigators to demonstrate independent convergence of numerous behavioral, developmental, and physiological traits (Chin et al., 2018; Kowalko, 2020; Riddle et al., 2018; Stockdale et al., 2018; Varatharasan et al., 2009). Our goal is to apply a neurophysiological approach across multiple *A. mexicanus* populations to examine the functional evolution of the lateral line sensory system.

The mechanoreceptive hair cells of the lateral line system detects fluid motion relative to the body and play an important role in essential behaviors (McHenry et al., 2009; Mekdara et al., 2018; Olszewski et al., 2012; Oteiza et al., 2017; Stewart et al., 2013). Cavefish have evolved anatomical enhancements of the lateral line, presumably to compensate for the loss of vision (Kowalko, 2020; McGaugh et al., 2020; Teyke, 1990; Yoshizawa et al., 2014). These anatomical alterations have been linked to substantial changes in behavior (Lloyd et al., 2018; Yoshizawa et al., 2010). However, almost nothing is known about physiological changes that accompany evolution, despite the fact that the response of peripheral senses to environmental change has been well documented (Kelley et al., 2018; McBride, 2007).

Endogenous depolarizations within sensory cells are transmitted to afferent neurons (hereafter “afferents”) as spontaneous action potentials (hereafter “spikes”), and are essential for maintaining a state of responsiveness and sensitivity (Dey et al., 2021; Douglass et al., 1993; Köppl, 1997) (Kiang, 1965; Manley & Robertson, 1976). This is true in the lateral line, where spontaneous depolarizing currents within the hair cell maintain a resting potential within the critical activation range of channels. This range is required to ensure transmitter release; thus these currents decrease the detection threshold of the system (Trapani & Nicolson, 2011). Spontaneous afferent activity is an established and reliable target for probing the neurophysiological basis of sensitivity across taxa (Hedwig, 2006; Krasne & Bryan, 1973; Mohr et al., 2003). Here, we use spontaneous afferent activity as an entry point into understanding the neural mechanism underlying lateral line function in cavefish.

Another important mechanism that determines lateral line sensitivity is an inhibitory feedback effect of the efferent system during swimming. Feedback mechanisms in general sculpt sensory systems in important ways; for example, by changing detection thresholds by altering the transmission frequency of afferent spikes (Crapse & Sommer, 2008; Straka et al., 2018). The efferent system of hair cells in particular has repeatedly evolved to modulate sensory processing (Köppl, 2011). More specifically, hindbrain efferent neurons (hereafter “efferents”) issue predictive signals that transmit in parallel to locomotor commands, termed corollary discharge (CD). CDs inhibit afferent activity to mitigate sensor fatigue that can result from self-generated stimuli (Russell & Roberts, 1972). CD is an important mechanism for sensitivity enhancement but has rarely been implicated for ecologically-relevant behaviors. For example, active-flow sensing by cavefish depends on detecting reafferent signals while swimming (Tan et al., 2011). Increased reliance on self-generated fluid motion (Odstrcil et al., 2021; Patton et al., 2010; Teyke, 1985) is divergent from our current understanding of the CD’s role in predictive motor signaling in the lateral line (Lunsford et al., 2019; Pichler & Lagnado, 2020).

For the first time, we identify a neurophysiological mechanism that has convergently evolved across *A. mexicanus* populations to increase hair cell sensitivity after eye loss. By investigating how differences in afferent and efferent signaling contribute to sensory enhancement in a comparative model, we provide insight into a potentially ubiquitous mechanism for sensory evolution.

## Results

Neuromasts of surface fish and Pachón cavefish larvae (6 days post fertilization; dpf) were labeled via 2-[4- (Dimethylamino)styryl]-1-ethylpyridinium iodide (DASPEI) staining and subsequently imaged (Figure 1A-B).Anterior lateral line neuromasts had previously been shown to differ in quantities and morphology as early as 2 months post fertilization between surface and cave fish (Yoshizawa et al., 2010). Here we show that Pachón cavefish exhibit this significant increase in anterior neuromast quantity as early as 6 dpf (p < 0.01, t = 3.168, df = 29; Figure 1C).

**Figure 1.**
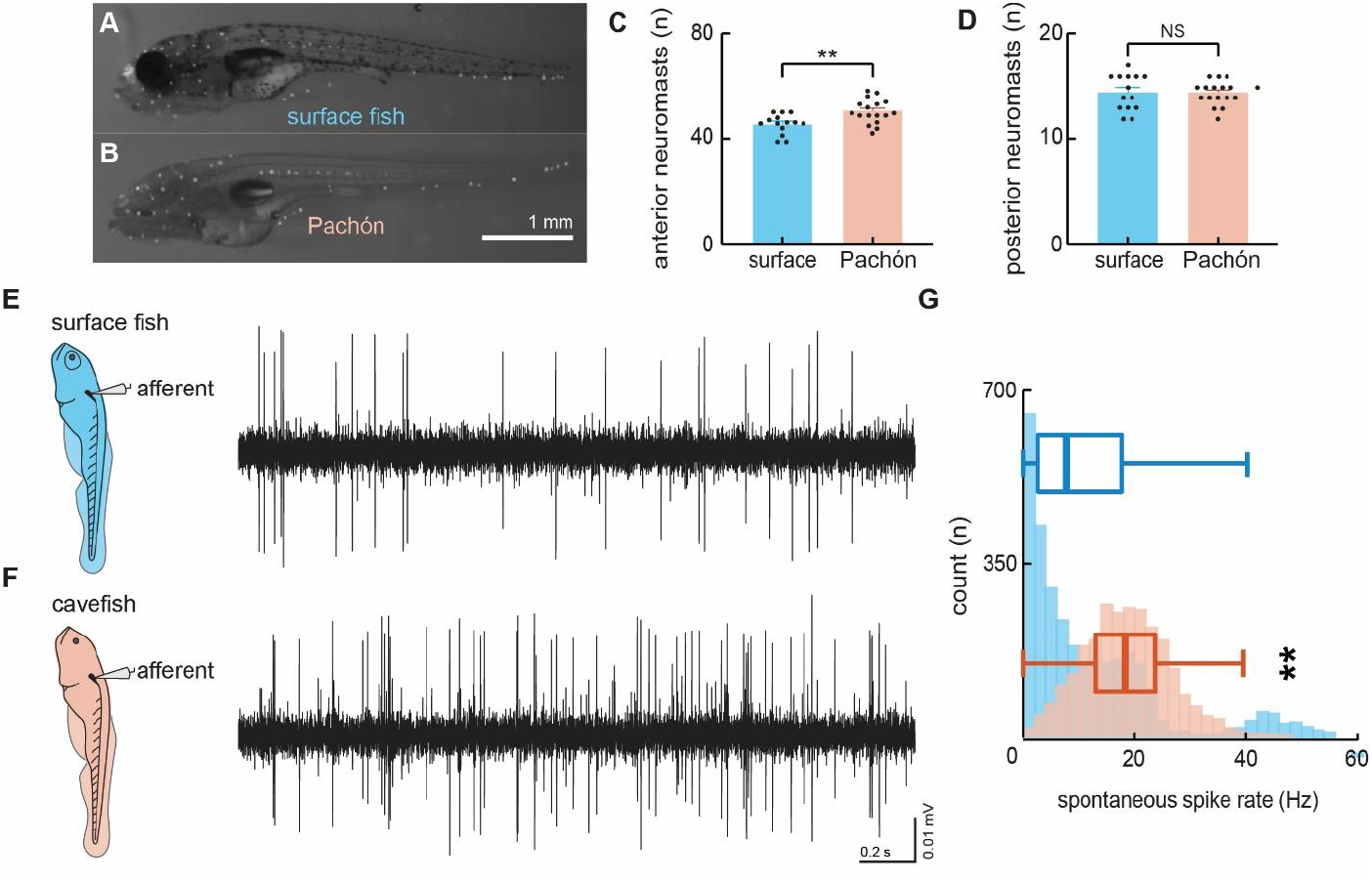
Spontaneous afferent neuron activity is elevated in blind cavefish. DASPEI staining of 6 dpf surface (**A**) and cavefish (**B**) show significantly different quantities of anterior lateral line neuromasts (**C**) and similar quantities of posterior neuromasts (**D**). Extracellular recordings were made in posterior lateral line afferents where the neuromast densities were similar in order to resolve the differences observed in afferent activity between larval surface fish (**E**) and Pachón cavefish (**F**) between 4-7 dpf. Number of occurrences and median intrinsic spike rates in both surface (blue; 12.4 Hz, n=10 fish) and Pachón (red; 18.6 Hz, n=5 fish) fish suggests that lateral line response thresholds in cavefish are lower than those of surface fish (**G).** Error bars are ± SE.

In contrast, both populations exhibit a similar number of posterior lateral line neuromasts (p = 0.77, U = 111.5; Figure 1D). Therefore, to investigate differences in underlying neurophysiology, we concentrated on the posterior lateral line system where neuromast density is similar across blind and surface morphs. We exclusively probed the posterior lateral line afferent neurons to establish whether sensory systems that are anatomically similar exhibit neurophysiological differences that contribute to enhanced sensitivity.

To examine the physiological basis of differences in lateral line function across surface and cavefish we used extracellular lateral line recordings adapted from protocols used in zebrafish (Figure 1E-F). Extracellular recordings of posterior lateral line afferents revealed intrinsic spontaneous activity was higher in Pachón cavefish (18.6 ± 0.2 Hz) while the animal was at rest, relative to surface fish (12.4 ± 0.3 Hz; p < 0.01, t = 15.97, df = 5,795; Figure 1G). Instantaneous afferent spike rate demonstrates substantial decreases during swimming in surface fish (surface = 3,167 swim bouts) and little effect in Pachón cavefish (Pachón = 2,612 swim bouts; Figure 2 A-B). We quantified and compared spike rates during swimming relative to the pre-swim period to examine patterns of the inhibitory effect between populations. During most surface fish swim bouts there was a reduction in afferent activity (n = 1,966/2,291, 85.8%), many of which resulted in complete quiescence of transmissions (n = 1,112/2,291, 48.5%). Conversely, afferent activity partially reduced during many swim bouts in Pachón cavefish (n = 1,303/2,439, 53.4%), but very few instances led to complete inhibition (n = 275/2,439, 11.3%). The distributions of relative spike rates during swimming reveal surface fish have a higher likelihood of experiencing no afferent activity during swimming while cavefish experience afferent activity during swimming similar to that of pre-swim activity levels (Figure 2 C). Therefore, surface fish experience significantly higher levels of inhibition (68.5 ± 0.01%) compared to cavefish (28.9 ± 0.01%; p < 0.01, t = 36.5, df = 4449; Figure 2 D). Surface fish demonstrate lateral line inhibition during swimming comparable to other fishes with intact visual systems (Flock & Russell, 1973; Lunsford et al., 2019; Pichler & Lagnado, 2020; Russell & Roberts, 1972) suggesting cavefish have evolved a unique functional phenotype for sensory gain control.

**Figure 2.**
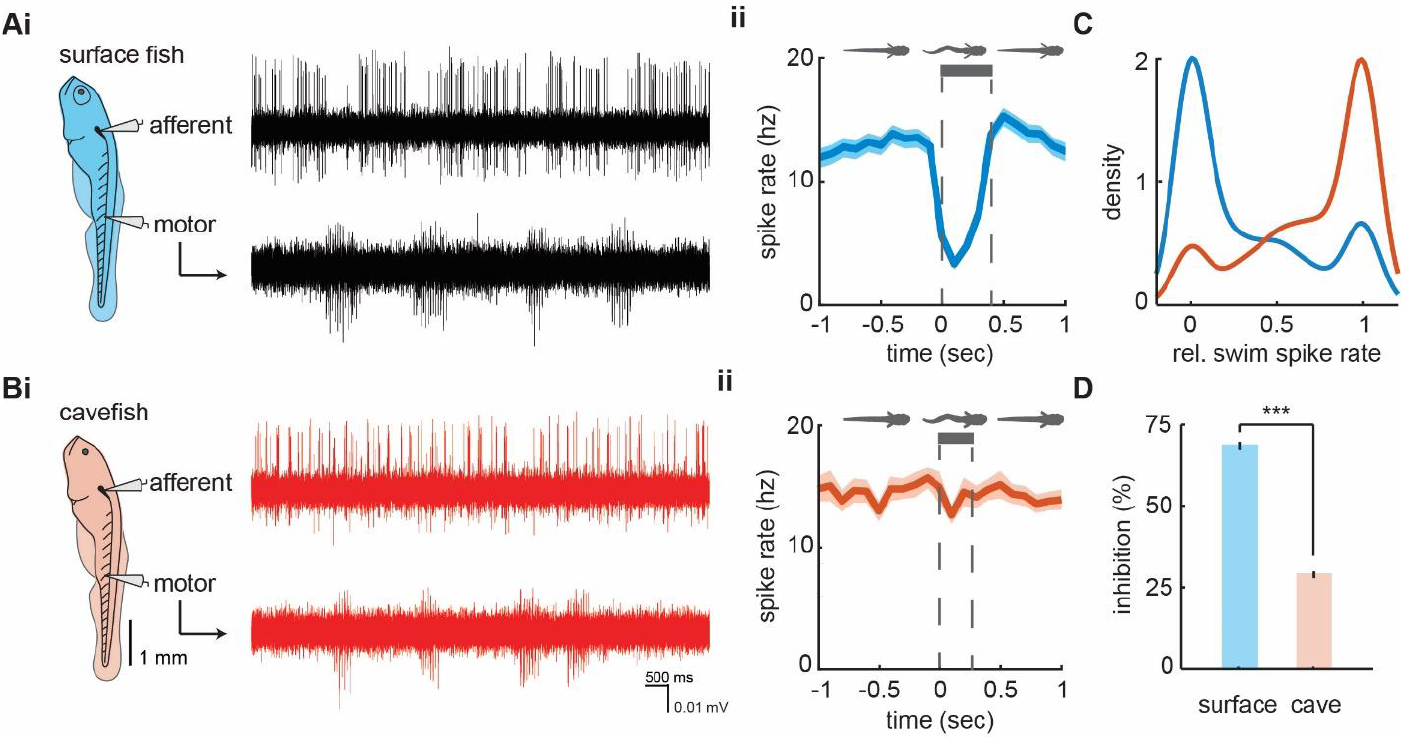
Afferent spike rate decreases during swimming in surface fish but not cavefish. Simultaneous recordings from afferent neurons from the posterior lateral line afferent ganglion and ventral motor roots along the body in (**Ai**) surface fish and (**Bi**) Pachón cavefish. Afferent spike rate decreases at the onset of swimming (time = 0) in (**Aii**) surface fish while (**Bii**) Pachón cavefish spike rate remained relatively constant during swimming. Bars represent average swim duration for surface fish (0.39 sec, n = 2,272 swims) and Pachón cavefish (0.27 sec, n = 2,429 swims). **C.** Kernel density estimate of spike rate during swimming relative to the pre-swim interval in both surface fish (blue) and cavefish (red). **D.** Surface fish experience greater levels of inhibition during swimming than cavefish. Significance level indicated by ‘***’. Error bars are ± SE.

We imaged hindbrain cholinergic efferent neurons to determine anatomical and functional connectivity. Between populations, backfilled efferents revealed similar soma quantities (surface: 2.4 ± 0.3; cave: 3.2 ± 0.5; p = 0.2, t = 1.36, df = 18) and size (surface: 62.1 ± 3.6 μm^2^; cave: 70.9 ± 4.5 μm^2^; p = 0.1, t = 1.50, df = 51; Figure 3 A-C). From electrophysiological recordings, we observed average spike rates during and prior to a swim were not positively correlated in control surface fish (r^2^ = 0.15, F1,13 = 2.3, p = 0.2), but the slope of the line indicates a fractional suppression of 78% that is significantly less than unity (slope 0.1, confidence interval (CI): −0.2-0.4; Figure 3 Di). After efferent ablation, surface fish average spike rates during and prior to a swim were not distinguishable from unity (slope 1.2, CI = −0.1-2.4) indicating that spike rates during swimming intervals differ from non-swimming intervals no differently than chance and therefore there is no detectable inhibitory effect. While swimming, surface fish without functioning efferent neurons demonstrated a signaling phenotype similar to both intact cavefish (slope 0.88, CI = −0.5-2.2) and cavefish with ablated efferents (slope 0.9, CI = 0.1-1.8; Figure 3 Dii). Therefore, putative cholinergic hindbrain efferents are necessary for afferent inhibition in surface fish, but do not demonstrate modulatory control of afferents in cavefish.

**Figure 3.**
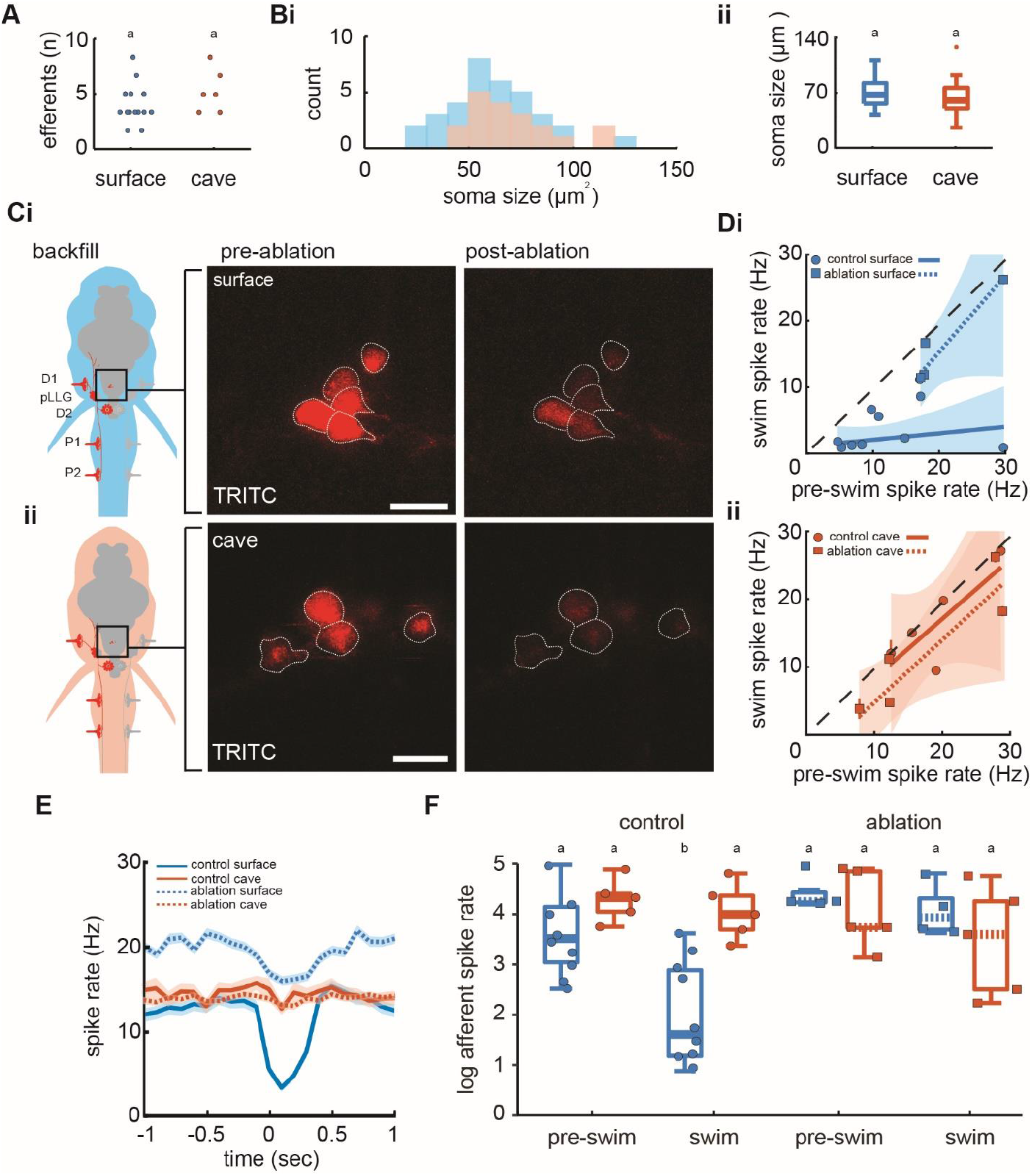
Efferent neurons are necessary for inhibition observed in afferents during swimming in surface fish but not cavefish. **A.** Backfilled hindbrain cholinergic efferent neurons were present in comparable numbers (2-3 cells) in both surface fish (n = 14) and cavefish (n = 6). **Bi.** Efferent soma size in surface (blue) and cavefish (red) is similar in both populations (**ii**). **C.** Efferent cell bodies were identified by backfilling rhodamine through posterior lateral line neuromasts in both surface fish (**i**) and cavefish (**ii**) and were ablated with a 30-s UV pulse. Scale bar: 20 μm. **D.** The line of best fit of spike rates before compared to during the swim significantly excludes unity in non-ablated, control surface fish (circle), but not in ablated surface fish (square), implying spike rate suppression in the former but not the latter (**i**). The slope of the line for control fish suggests the inhibition is not correlated to the spontaneous afferent activity preceding the swim. The line of best fit of Pachón cavefish pre-swim and swim spike rates did not exclude unity in both control (circle) and ablated (square) treatments (**ii**). Dashed line indicates the line of unity, corresponding to no average difference of spike rate during swimming. **E.** Instantaneous spike rates of Pachón cavefish were unaffected by ablating the lateral line whereas the inhibitory effect was eliminated in ablated surface fish. Time is relative to the onset of motor activity. **F.** Surface fish (blue) display reduced spike rates during swimming compared to before swimming in control fish. Pachón cavefish (red) did not display reduced spike rate during swimming in neither control nor ablated treatments. Ablated surface individuals also did not display reduced spike rate during swimming resulting in a signaling phenotype comparable to cavefish. Groupings of statistical similarity are denoted by ‘a’ and ‘b’, whereas a is significantly different from b. All error bars represent ± SE.

We compared pre-swim and swim spike rates across populations and treatments to determine efferent contribution to inhibition (Figure 3 E). We found significant differences in afferent activity among pre-swim and swim intervals (F_7,40_ = 7.6, p < 0.01; Figure 3 F). Surface fish afferent spike rates during swimming (3.9 ± 0.1 Hz) were 71% lower than the immediate pre-swim period in control fish (13.7 ± 0.2 Hz; Tukey’s post hoc test, p < 0.01). Post-swim spike rate (13. 7 ± 0.3 Hz) recovered to pre-swim spike rate. In control Pachón cavefish, we observed some decrease in afferent spike rate during swimming (16.8 ± 0.2 Hz) but it was not statistically discriminated from pre-swim (20.5 ± 0.2 Hz; Tukey’s post hoc test, p = 0.94). Efferent ablation in surface fish resulted in afferent activity during swimming (17.4 ± 0.3 Hz) to increase to pre-swim levels (20.7 ± 0.3 Hz; Tukey’s post hoc test, p = 0.99). Ablated surface fish also demonstrated spike dynamics during swimming comparable to ablated and control cavefish (Pachón ablated, pre-swim: 18.6 ± 0.3 Hz; Pachón ablated, swimming: 14.7 ± 0.3 Hz). These findings indicate that efferents are necessary for inhibition of afferents during swimming in surface fish, but not cavefish.

We compared lateral line activity between three different cave populations (Figure 4 A); the Tinaja and Pachón populations, which are derived from similarly timed colonization events, and the Molino population that is derived from a more recent colonization event (Bradic et al., 2012; Dowling et al., 2002; Herman et al., 2018). Blind cavefish populations exhibited similar spontaneous spike rates across populations (F_2,17_ = 0.68, p = 0.52), and swimming showed little effect on lateral line activity across blind cavefish populations (Figure 4 B). We observed a minor decrease in afferent spike rates across cavefish populations and the relative change (i.e. inhibition) was similar in Pachón and Tinaja (Tukey’s post hoc test; Pachón: 0.28 ± 0.01; Tinaja: 0.32 ± 0.02; Figure 4 C). Molino demonstrated an intermediate phenotype compared to Pachón and Tinaja (Tukey’s post hoc test; Molino: 0.23 ± 0.01; p < 0.01). The Molino population (new lineage) is most distantly related to the Pachón and Tinaja populations (old lineage), originating from a more recent surface fish colonization of caves thus providing phylogenetic evidence that coincides with statistically similar groupings (Herman et al., 2018). We examined the correlation between non-swimming and swimming spike rates across cave populations to determine whether the inhibitory effect was significant. Spike rates during swimming and pre-swim intervals were not positively correlated in all cave populations, and afferent spike rates during swimming were not statistically distinguishable from unity across populations (Pachón: slope = 0.88, CI = −0.5 – 2.2; Tinaja: slope = 1.03, CI = −0.2 - 2.2; Molino: slope = 1.02, CI = 0.8 - 1.2; Figure 4 D), indicating there was no significant decrease in afferent activity during swimming within blind cavefish populations.

**Figure 4.**
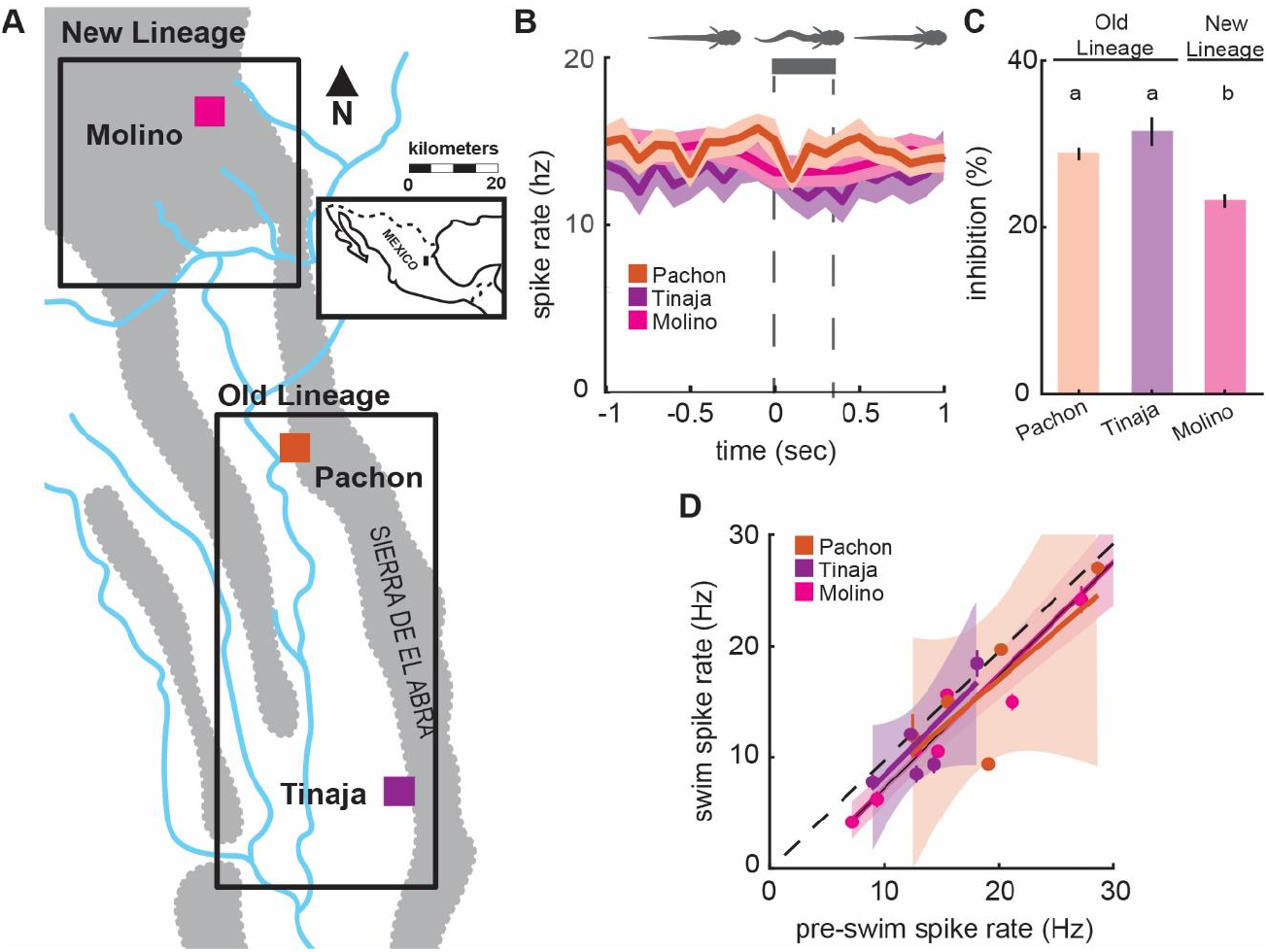
Enhanced lateral line sensitivity during swimming convergently evolved across three blind populations. **A.** Molino cave populations (pink; New Lineage) have evolved more recently relative to Pachón (red) and Tinaja (purple; Old Lineage) cave populations. Lineage delineations inferred from phylogenetic data (Herman et al. 2018). **B.** Mean spontaneous afferent spike rate remains constant at the onset of fictive swimming (time = 0) in Pachón (n = 5), Molino (n = 8), and Tinaja (n = 5) populations. Bar represents average swim duration for Pachón (0.27 sec, n = 2,429 swim bouts), Molino (0.42 sec, n = 1,474 swim bouts), and Tinaja (0.35 sec, n = 464). **C.** Percent change in spike rate from pre-swim to swim intervals (i.e. inhibition) was small, but significantly different between blind cave populations. Post-hoc Tukey test revealed that Molino cavefish experienced significantly less reduction in spike rate when compared to Pachón and Tinaja populations. Statistically similar groups are indicated by ‘a’ and ‘b’. **D.** The line of best fit of pre-swim and swim spike rates does not significantly exclude unity in any of the blind cavefish populations implying there is no detectable inhibitory effect. Dashed line indicates the line of unity, corresponding to no average difference of spike rate during swimming. All values represent mean ± SE.

## Discussion

Our principle finding indicates that elevated afferent activity underlies the increased responsiveness of cavefish to flow stimuli. The increased afferent activity in cavefish is a likely consequence of eye-loss, which has robust effects on other physiological systems (Duboué et al., 2011; Varatharasan et al., 2009). Heightened lateral line sensitivity in adults has been previously attributed to increased neuromast density in the anterior lateral line (ALL) (J. Jaggard et al., 2017; Lloyd et al., 2018; McHenry et al., 2008; Patton et al., 2010; Teyke, 1990; Yoffe et al., 2020, 2020; Yoshizawa et al., 2010). This difference in neuromast quantity does not exist in the posterior lateral line at the larval stage, allowing us a unique opportunity to investigate the neural mechanisms that can enhance flow sensing in a strong model for evolution. Our discovery of increased spontaneous activity in cavefish is a powerful addition for flow sensing, and we predict that this in combination with the increased density of neuromasts in the ALL is what ultimately enables cavefish to perform active flow sensing (Yoshizawa et al., 2010).

Higher spontaneous activity is one of two mechanisms that are responsible for higher lateral line sensitivity. The other involves the inhibitory effects of the efferent system during swimming, a feedback mechanism that is conserved across the diversity of fishes (Flock & Russell, 1973; Lunsford et al., 2019; J. Montgomery et al., 1996; J. C. Montgomery & Bodznick, 1994; Roberts & Russell, 1972; Tricas & Highstein, 1991). This is true in swimming surface fish but not in cavefish. We found that three blind populations of cavefish (i.e. Pachón, Molino, Tinaja) have repeatedly lost the capability for efferent modulation during swimming. When one considers that lateral line efferent activity can be driven by direct inputs from the visual system in zebrafish (Reinig et al., 2017), it seems possible that eye degeneration induces the loss of efferent function. This interpretation is consistent with the idea that lateral line efferents are thought to have undergone regressive loss before in the ancestral lamprey and hagfish (Kishida et al., 1987; Köppl, 2011; Koyama et al., 1990), both of which are nearly or completely blind during development (Dickson & Collard, 1979; Fernholm & Holmberg, 1975). However, our results illustrate that cholinergic efferent system is still present in cavefish, having lost functionality rather than disappearing (the efferent system is functional in surface fish). Exploring pre- and postsynaptic differences such as acetylcholine release or the density of nicotinic acetylcholine receptors (nAChR) may explain the reduced inhibitory efficacy and reveal molecular targets that could disrupt efferent function over the course of evolution (Dawkins et al., 2005).

Our demonstration of CD inactivity in cavefish provides an alternative mechanism by which evolution can enhance sensitivity, one that proceeds by decreasing inhibition rather than augmenting sensor morphology or density (Yamamoto et al., 2009; Yoshizawa et al., 2010, 2014). The impact of increasing sensitivity through a lack of inhibition is apparent during active-flow sensing in adult *A. mexicanus*. Active-flow sensing occurs during swimming and involves using the reflection of self-generated flow fields (Bleckmann et al., 1991) to follow walls (Patton et al., 2010), avoid obstacles (Teyke, 1985; Windsor et al., 2008), and discrimination between shapes (De Perera, 2004; Hassan, 1989; Von Campenhausen et al., 1981). The repeated loss of inhibitory feedback across blind cavefish populations suggests that it is easier to cease function than to develop more neuromasts or other additive alternatives.

Cavefish swim by using different body motions than surface fish. This finding has been interpreted as a mechanism to enhance wall following, which occurs when the bow wake of a swimming cavefish is reflected off of a solid surface and then detected (Patton et al., 2010; Sharma et al., 2009). Altered swimming kinematics is also thought to have arisen from a general increase in sensitivity to flow (Tan et al., 2011; Windsor et al., 2008). Our results provide an alternate suggestion; the lack of efferent function in cavefish precludes the sensory feedback necessary for sensing self-movement and body position in water (proprioception). Corollary discharge, a parallel motor command that decreases the afferent activity of fishes when swimming, has recently been found to play a critical role in swimming efficiency by enabling the tracking of the traveling body wave during undulation (Skandalis et al., 2021). We hypothesize that the evolved loss of efferent function that enables cavefish to successfully avoid collisions in subterranean habitats is likely favored over optimizing swimming efficiency (Nakamura, 1997; Uysal et al., 2010). We predict that loss of efferent function will be found in other blind hypogean species (Costa Sampaio et al., 2012) and that their respective surface populations will possess intact efferent functionality, as we have found in *A. mexicanus*. Neurophysiological recordings across a wider diversity of species would provide valuable insight into how efferent function may be sculpted by environmental selection and phylogenetic membership.

By employing neurophysiological approaches in the lateral line system of *A. mexicanus* for the first time, we show that elevated lateral line afferent activity and loss of efferent function have repeatedly evolved together across cavefish populations. Our findings come at a time when genetic tools in *A. mexicanus* enable brain-wide imaging and gene-editing based screening to identify candidate neural circuits and genes critical in the evolution of sensory systems (Jaggard et al., 2020; Warren et al., 2021). Going forward, applying genetic and electrophysiological tools in well-characterized neural circuits promises to inform our understanding of the evolution of neural systems and behavior more broadly.

## Materials and Methods

### Animals

Fish were progeny of pure-bred stocks originally collected in Mexico (Duboué et al., 2011) that have been maintained at the Florida Atlantic University core facilities. Larvae were raised in 10% Hank’s solution (137 mM NaCl, 5.4 mM KCl, 0.25 mM Na_2_HPO_4_, 0.44 mM KH_2_PO_4_, 1.3 mM CaCl_2_, 1.0 mM MgSO_4_, 4.2 mM NaHCO_4_; pH 7.3) at 26°C. All experiments were performed according to protocols approved by the University of Florida or Florida Atlantic University Institutional Animal Care and Use Committee. Animal health was assessed by monitoring blood flow throughout each experiment.

### Neuromast imaging

To assess neuromast quantities, larvae aged six dpf were submerged in 5 μg/ml DASPEI dissolved in embryo medium for 15 minutes as previously described (Van Trump et al., 2010). Larvae were then transferred to ice-cold water for 30-45 seconds then immersed in 8% methylcellulose for imaging. Images were taken using a Nikon DS-Qi2 monochrome microscope camera mounted on a Nikon SMZ25 Stereo microscope (Nikon; Tokyo, Japan). Neuromasts innervated by posterior lateral line afferents and anterior lateral line afferents were tabulated separately.

### Electrophysiology

Prior to recordings, *A. mexicanus* larvae (4-7 dpf) were paralyzed using 0.1% α-bungarotoxin (Lunsford & Liao, 2021). Once paralyzed, larvae were then transferred into extracellular solution (134 mM NaCl, 2.9 mM KCl, 1.2 mM MgCl_2_, 2.1 mM CaCl_2_, 10 mM glucose, 10 mM HEPES buffer; pH 7.8, adjusted with NaOH) and pinned with etched tungsten pins through their dorsal notochord and otic vesicle into a Sylgard-bottom dish.

Multiunit extracellular recordings of the posterior lateral line afferent ganglion were made in surface fish (n = 10) and Pachón cave fish (n = 5). Prior to recording from the afferent neurons, a bore pipette was used to break through the skin to expose the afferent soma. Recording electrodes (~30 μm tip diameter) were pulled from borosilicate glass (model G150F-3, inner diameter: 0.86, outer diameter: 1.50; Warner Instruments, Hamden, CT) on a model P-97 Flaming/Brown micropipette puller (Sutter Instruments, Novato, CA) and filled with extracellular solution. Once contact with afferent somata was achieved, gentle negative pressure was applied (20-50 mmHg; pneumatic transducer tester, model DPM1B, Fluke Biomedical Instruments, Everett, WA). Pressure was adjusted to atmospheric (0 mmHg) once a stable recording was achieved. Simultaneously, ventral root (VR) recordings were made through the skin (Masino & Fetcho, 2005) to detect voluntary fictive swimming. All recordings were sampled at 20 kHz and amplified with a gain of 1000 in Axoclamp 700B, digitized with Digidata 1440A and saved in pClamp10 (Molecular Devices).

All recordings were analyzed in Matlab (vR2016b) using custom written scripts. Both spontaneous afferent spikes and swimming motor activity identified using a combination of spike parameters previously described (Lunsford et al., 2019). Afferent neuron activity within a time interval equal to the subsequent fictive swim bout, hereafter termed “pre-swim”, was quantified and compared to afferent activity during swimming to measure relative changes in spontaneous firing.

### Efferent Ablations

Hindbrain efferent neurons were backfilled with tetramethylrhodamine (TRITC, 3 kDa; Molecular Probes, Eugene, OR). *A. mexicanus* larvae (4 dpf) were anesthetized in MS-222 (Tricaine, Western Chemical Inc. Ferndale, WA) and embedded in agar. To selectively label the hindbrain cholinergic efferent neurons, we systematically electroporated (Axoporator 800A Single Cell Electroporator, Axon CNS Systems, Molecular Devices LLC, San Jose, CA) TRITC into the efferent terminals that innervate the D1, D2, L1, and L2 neuromasts of the lateral line in surface fish (n = 14) and Pachón cavefish (n = 7). Electroporation does not ensure labelling of all efferent neurons so we standardized parameters (30 V, 50 Hz, 500 ms, square pulse) and targeted the same neuromasts across populations to minimize variation in labelling efficacy. Larvae were then gently freed from the agar and allowed to swim freely and recover overnight. Larvae (5 dpf) were then paralyzed via α-bungarotoxin immersion, remounted in agar dorsal surface down, and imaged on a Leica SP5 confocal microscope (Leica Microsystems, Wetzlar, Germany). Efferent soma size and quantity was measured within identified TRITC-labelled cells in ImageJ (v1.48; U. S. National Institutes of Health, Bethesda, MD). To perform targeted ablations of surface fish (n = 5) and cavefish (n = 6) efferent neurons, a near-ultraviolet laser was focused at a depth corresponding to the maximum fluorescence intensity of each soma, to ensure we were targeting its centre. We applied the FRAP Wizard tool in Leica application software to target individual cells. We ablated target cells with a 30 s exposure to the near-ultraviolet laser line (458 nm), and successful targeting was confirmed by quenching of the backfilled dye. This method has been successfully applied and validated in similar systems (Liu & Fetcho, 1999; Soustelle et al., 2008). Fish were again freed from agar and allowed to swim freely and recover overnight. Electrophysiological recordings were performed on ablated surface fish (n = 4) and cavefish (n = 5) at 6 dpf to simultaneously monitor afferent activity and motor activity.

### Statistical analysis

Neuromast data were analyzed using GraphPad Prism 8.4.3. Normality was assessed via Shapiro-Wilk test. Anterior lateral line neuromast count data were found to be normally distributed. Anterior lateral line neuromast quantities in surface and cave larvae were compared using an unpaired t-test. Posterior lateral line neuromast count was found to not be normally distributed and was subsequently analyzed via Mann-Whitney U-test

Analyses of electrophysiological data were performed using custom written models in the R language (R development core team, vR2016b) using packages car, visreg, reshape2, plyr, dplyr, ggplot2, gridExtra, minpack.lm, nlstools, investr, and cowplot (Auguie et al., 2017; Baty et al., 2015; Breheny & Burchett, 2017; Fox & Weisberg, 2018; Wickham, 2007, 2009; Wickham et al., 2019; Wilke, 2019). Spontaneous afferent spike rate was calculated by taking the number of spikes over the duration of time where the larva was inactive. Instantaneous afferent spike rate was calculated using a moving average filter and a 100 ms sampling window. Pre-swim and swim spike rate were calculated by taking the number of spikes within the respective period over its duration. Pre-swim periods of inactivity made it challenging to interpret changes in afferent activity, so we restricted the dataset to only include swim bouts that were preceded by a minimum of one afferent spike within the pre-swim interval (surface = 2,291 swim bouts; Pachón = 2,429 swim bouts; Molino = 1.474 swim bouts; Tinaja = 464 swim bouts). Swim frequency was calculated by taking the number of bursts within a swim bout over the duration of the swim bout. Relative spike rate was calculated by taking the swim spike rate over the pre-swim spike rate. All variables were averaged for each individual fish. The precision of estimates for each individual is a function of the number of swims, so we analyzed variable relationships using weighted regressions, with individual weights equal to the square root of the number of swims. We log transformed variables in which the mean and the variance were correlated. To quantify the inhibition of the afferent spike frequency during swimming we tested for a significant difference in afferent spike frequency during swimming as compared to non-swimming periods using a paired sample student’s t-test.

Differences in afferent spike rates between the periods of interest (pre-swim and swim) across populations and treatments were tested by N-way analysis of variance (ANOVA) followed by Tukey’s post-hoc test to detect significant differences in spike rates between swim periods or treatments. Linear models were used to detect relationships between spike rate during swimming and other independent variables (e.g. spike rate pre-swim). Data is shown throughout the manuscript as mean ± standard error. Statistical significance is reported at α = 0.05.

## Acknowledgements

We gratefully acknowledge support from the US-Israel BSF SP#2018-190, National Science Foundation (IOS165674), and National Institute of Health (1R01GM127872) to ACK, and the National Institute of Health (DC010809), National Science Foundation (IOS1856237, IOS2102891), and support from the Whitney Laboratory for Marine Biosciences to JCL.

## Author Contributions

E.T.L. and J.C.L conception and design of research; E.T.L. performed electrophysiology and hindbrain imaging experiments; A.P. performed lateral line imaging; E.T.L. and A.P. analyzed data; E.T.L, A.P., A.C.K. and J.C.L. interpreted results of experiments; E.T.L. prepared figures; E.T.L. drafted manuscript; E.T.L., A.P., A.C.K. and J.C.L. edited and revised manuscript; E.T.L., A.P., A.C.K. and J.C.L. approved final version of manuscript.

## References

Auguie, B., Antonov, A., & Auguie, M. B. (2017). Package ‘gridExtra.’ Miscellaneous Functions for “Grid” Graphics.

Baty, F., Ritz, C., Charles, S., Brutsche, M., Flandrois, J.-P., & Delignette-Muller, M.-L. (2015). A toolbox for nonlinear regression in R: The package nlstools. Journal of Statistical Software, 66(5), 1–21.

Bleckmann, H., Breithaupt, T., Blickhan, R., & Tautz, J. (1991). The time course and frequency content of hydrodynamic events caused by moving fish, frogs, and crustaceans. Journal of Comparative Physiology A, 168(6), 749–757.

Blin, M., Tine, E., Meister, L., Elipot, Y., Bibliowicz, J., Espinasa, L., & Rétaux, S. (2018). Developmental evolution and developmental plasticity of the olfactory epithelium and olfactory skills in Mexican cavefish. Developmental Biology, 441(2), 242–251.

Bradic, M., Beerli, P., García-de León, F. J., Esquivel-Bobadilla, S., & Borowsky, R. L. (2012). Gene flow and population structure in the Mexican blind cavefish complex (Astyanax mexicanus). BMC Evolutionary Biology, 12(1), 9. https://doi.org/10.1186/1471-2148-12-9

Breheny, P., & Burchett, W. (2017). Visualization of regression models using visreg. R J., 9(2), 56.

Chin, J. S. R., Gassant, C. E., Amaral, P. M., Lloyd, E., Stahl, B. A., Jaggard, J. B., Keene, A. C., & Duboue, E. R. (2018). Convergence on reduced stress behavior in the Mexican blind cavefish. Developmental Biology, 441(2), 319–327. https://doi.org/10.1016/j.ydbio.2018.05.009

Costa Sampaio, F. A., Pompeu, P. S., de Andrade e Santos, H., & Lopes Ferreira, R. (2012). Swimming performance of epigeal and hypogeal species of Characidae, with an emphasis on the troglobiotic Stygichthys typhlops Brittan & Böhlke, 1965. International Journal of Speleology, 41(1), 2.

Crapse, T. B., & Sommer, M. A. (2008). Corollary discharge across the animal kingdom. Nature Reviews. Neuroscience, 9(8), 587–600. https://doi.org/10.1038/nrn2457

Dawkins, R., Keller, S. L., & Sewell, W. F. (2005). Pharmacology of acetylcholine-mediated cell signaling in the lateral line organ following efferent stimulation. Journal of Neurophysiology, 93(5), 2541–2551. https://doi.org/10.1152/jn.01283.2004

De Perera, T. B. (2004). Fish can encode order in their spatial map. Proceedings of the Royal Society of London. Series B: Biological Sciences, 271(1553), 2131–2134.

Dey, A., Zele, A. J., Feigl, B., & Adhikari, P. (2021). Threshold vision under full-field stimulation: Revisiting the minimum number of quanta necessary to evoke a visual sensation. Vision Research, 180, 1–10.

Dickson, D. H., & Collard, T. R. (1979). Retinal development in the lamprey (Petromyzon marinus L.): Premetamorphic ammocoete eye. American Journal of Anatomy, 154(3), 321–336.

Douglass, J. K., Wilkens, L., Pantazelou, E., & Moss, F. (1993). Noise enhancement of information transfer in crayfish mechanoreceptors by stochastic resonance. Nature, 365(6444), 337–340.

Dowling, T. E., Martasian, D. P., & Jeffery, W. R. (2002). Evidence for multiple genetic forms with similar eyeless phenotypes in the blind cavefish, Astyanax mexicanus. Molecular Biology and Evolution, 19(4), 446–455.

Duboué, E. R., Keene, A. C., & Borowsky, R. L. (2011). Evolutionary convergence on sleep loss in cavefish populations. Current Biology, 21(8), 671–676. https://doi.org/10.1016/j.cub.2011.03.020

Fernholm, B., & Holmberg, K. (1975). The eyes in three genera of hagfish (Eptatretus, Paramyxine andMyxine)—A case of degenerative evolution. Vision Research, 15(2), 253–IN4.

Flock, A., & Russell, I. J. (1973). The post-synaptic action of efferent fibres in the lateral line organ of the burbot Lota lota. The Journal of Physiology, 235(3), 591–605.https://doi.org/10.1113/jphysiol.1973.sp010406

Fox, J., & Weisberg, S. (2018). An R companion to applied regression. Sage publications.

Hassan, E. S. (1989). Hydrodynamic imaging of the surroundings by the lateral line of the blind cave fish Anoptichthys jordani. In The mechanosensory lateral line (pp. 217–227). Springer.

Hedwig, B. (2006). Pulses, patterns and paths: Neurobiology of acoustic behaviour in crickets. Journal of Comparative Physiology A, 192(7), 677–689. https://doi.org/10.1007/s00359-006-0115-8

Herman, A., Brandvain, Y., Weagley, J., Jeffery, W. R., Keene, A. C., Kono, T. J., Bilandžija, H., Borowsky, R., Espinasa, L., & O’Quin, K. (2018). The role of gene flow in rapid and repeated evolution of cave-related traits in Mexican tetra, Astyanax mexicanus. Molecular Ecology, 27(22), 4397–4416.

Hoke, K., Schwartz, A., & Soares, D. (2012). Evolution of the fast start response in the cavefish Astyanax mexicanus. Behavioral Ecology and Sociobiology, 66(8), 1157–1164.

Hüppop, K. (1987). Food-finding ability in cave fish (Astyanax fasciatus). International Journal of Speleology, 16(1), 4.

Jaggard, J. B., Lloyd, E., Yuiska, A., Patch, A., Fily, Y., Kowalko, J. E., Appelbaum, L., Duboue, E. R., & Keene, A. C. (2020). Cavefish brain atlases reveal functional and anatomical convergence across independently evolved populations. Science Advances, 6(38), eaba3126.

Jaggard, J. B., Stahl, B. A., Lloyd, E., Prober, D. A., Duboue, E. R., & Keene, A. C. (2018). Hypocretin underlies the evolution of sleep loss in the Mexican cavefish. Elife, 7, e32637.

Jaggard, J., Robinson, B. G., Stahl, B. A., Oh, I., Masek, P., Yoshizawa, M., & Keene, A. C. (2017). The lateral line confers evolutionarily derived sleep loss in the Mexican cavefish. Journal of Experimental Biology, 220(2), 284–293. https://doi.org/10.1242/jeb.145128

Jourjine, N., & Hoekstra, H. E. (2021). Expanding evolutionary neuroscience: Insights from comparing variation in behavior. Neuron.

Kelley, J. L., Chapuis, L., Davies, W. I. L., & Collin, S. P. (2018). Sensory System Responses to Human-Induced Environmental Change. Frontiers in Ecology and Evolution, 6. https://doi.org/10.3389/fevo.2018.00095

Kiang, N. Y.-S. (1965). Discharge patterns of single fibers in the cat’s auditory nerve. MASSACHUSETTS INST OF TECH CAMBRIDGE RESEARCH LAB OF ELECTRONICS.

Kishida, R., Goris, R. C., Nishizawa, H., Koyama, H., Kadota, T., & Amemiya, F. (1987). Primary neurons of the lateral line nerves and their central projections in hagfishes. Journal of Comparative Neurology, 264(3), 303–310.

Köppl, C. (1997). Frequency tuning and spontaneous activity in the auditory nerve and cochlear nucleus magnocellularis of the barn owl Tyto alba. Journal of Neurophysiology, 77(1), 364–377. https://doi.org/10.1152/jn.1997.77.1.364

Köppl, C. (2011). Evolution of the octavolateral efferent system. In Auditory and vestibular efferents (pp. 217–259). Springer.

Kowalko, J. (2020). Utilizing the blind cavefish Astyanax mexicanus to understand the genetic basis of behavioral evolution. Journal of Experimental Biology, 223(jeb208835). https://doi.org/10.1242/jeb.208835

Koyama, H., Kishida, R., Goris, R. C., & Kusunoki, T. (1990). Organization of the primary projections of the lateral line nerves in the lamprey Lampetra japonica. Journal of Comparative Neurology, 295(2), 277–289.

Krasne, F. B., & Bryan, J. S. (1973). Habituation: Regulation through Presynaptic Inhibition. Science, 182(4112), 590–592.

Liu, K. S., & Fetcho, J. R. (1999). Laser ablations reveal functional relationships of segmental hindbrain neurons in zebrafish. Neuron, 23(2), 325–335.

Lloyd, E., Olive, C., Stahl, B. A., Jaggard, J. B., Amaral, P., Duboué, E. R., & Keene, A. C. (2018). Evolutionary shift towards lateral line dependent prey capture behavior in the blind Mexican cavefish. Developmental Biology, 441(2), 328–337. https://doi.org/10.1016/j.ydbio.2018.04.027

Lunsford, E. T., & Liao, J. C. (2021). Activity of Posterior Lateral Line Afferent Neurons during Swimming in Zebrafish. JoVE (Journal of Visualized Experiments), 168, e62233. https://doi.org/10.3791/62233

Lunsford, E. T., Skandalis, D. A., & Liao, J. C. (2019). Efferent modulation of spontaneous lateral line activity during and after zebrafish motor commands. Journal of Neurophysiology, 122(6), 2438–2448.

Manley, G. A., & Robertson, D. (1976). Analysis of spontaneous activity of auditory neurones in the spiral ganglion of the guinea-pig cochlea. The Journal of Physiology, 258(2), 323–336. https://doi.org/10.1113/jphysiol.1976.sp011422

Masino, M. A., & Fetcho, J. R. (2005). Fictive swimming motor patterns in wild type and mutant larval zebrafish. Journal of Neurophysiology, 93(6), 3177–3188. https://doi.org/10.1152/jn.01248.2004

McBride, C. S. (2007). Rapid evolution of smell and taste receptor genes during host specialization in Drosophila sechellia. Proceedings of the National Academy of Sciences, 104(12), 4996–5001. https://doi.org/10.1073/pnas.0608424104

McGaugh, S. E., Kowalko, J. E., Duboué, E., Lewis, P., Franz-Odendaal, T. A., Rohner, N., Gross, J. B., & Keene, A. C. (2020). Dark world rises: The emergence of cavefish as a model for the study of evolution, development, behavior, and disease. Journal of Experimental Zoology Part B: Molecular and Developmental Evolution, 334(7–8), 397–404. https://doi.org/10.1002/jez.b.22978

McHenry, M. J., Feitl, K. E., Strother, J. A., & Van Trump, W. J. (2009). Larval zebrafish rapidly sense the water flow of a predator’s strike. Biology Letters, 5(4), 477–479. https://doi.org/10.1098/rsbl.2009.0048

McHenry, M. J., Strother, J. A., & Van Netten, S. M. (2008). Mechanical filtering by the boundary layer and fluid–structure interaction in the superficial neuromast of the fish lateral line system. Journal of Comparative Physiology A, 194(9), 795.

Mekdara, P. J., Schwalbe, M. A. B., Coughlin, L. L., & Tytell, E. D. (2018). The effects of lateral line ablation and regeneration in schooling giant danios. The Journal of Experimental Biology, 221(Pt 8). https://doi.org/10.1242/jeb.175166

Mohr, C., Roberts, P. D., & Bell, C. C. (2003). The Mormyromast Region of the Mormyrid Electrosensory Lobe. I. Responses to Corollary Discharge and Electrosensory Stimuli. Journal of Neurophysiology, 90(2), 1193–1210. https://doi.org/10.1152/jn.00211.2003

Montgomery, J., Bodznick, D., & Halstead, M. (1996). Hindbrain signal processing in the lateral line system of the dwarf scorpionfish Scopeana papillosus. Journal of Experimental Biology, 199(4), 893–899.

Montgomery, J. C., & Bodznick, D. (1994). An adaptive filter that cancels self-induced noise in the electrosensory and lateral line mechanosensory systems of fish. Neuroscience Letters, 174(2), 145–148. https://doi.org/10.1016/0304-3940(94)90007-8

Nakamura, T. (1997). Quantitative analysis of gait in the visually impaired. Disability and Rehabilitation, 19(5), 194–197.

Odstrcil, I., Petkova, M. D., Haesemeyer, M., Boulanger-Weill, J., Nikitchenko, M., Gagnon, J. A., Oteiza, P., Schalek, R., Peleg, A., Portugues, R., Lichtman, J. W., & Engert, F. (2021). Functional and ultrastructural analysis of reafferent mechanosensation in larval zebrafish. Current Biology: CB, S0960-9822(21)01530-X. https://doi.org/10.1016/j.cub.2021.11.007

Olszewski, J., Haehnel, M., Taguchi, M., & Liao, J. C. (2012). Zebrafish larvae exhibit rheotaxis and can escape a continuous suction source using their lateral line. PloS One, 7(5), e36661. https://doi.org/10.1371/journal.pone.0036661

Oteiza, P., Odstrcil, I., Lauder, G., Portugues, R., & Engert, F. (2017). A novel mechanism for mechanosensory-based rheotaxis in larval zebrafish. Nature, 547(7664), 445–448.

Patton, P., Windsor, S., & Coombs, S. (2010). Active wall following by Mexican blind cavefish (Astyanax mexicanus). Journal of Comparative Physiology A, 196(11), 853–867.

Paz, A., McDole, B., Kowalko, J. E., Duboue, E. R., & Keene, A. C. (2020). Evolution of the acoustic startle response of Mexican cavefish. Journal of Experimental Zoology Part B: Molecular and Developmental Evolution, 334(7–8), 474–485.

Pichler, P., & Lagnado, L. (2020). Motor Behavior Selectively Inhibits Hair Cells Activated by Forward Motion in the Lateral Line of Zebrafish. Current Biology, 30(1), 150–157.e3. https://doi.org/10.1016/j.cub.2019.11.020

Reinig, S., Driever, W., & Arrenberg, A. B. (2017). The descending diencephalic dopamine system is tuned to sensory stimuli. Current Biology, 27(3), 318–333.

Riddle, M. R., Aspiras, A. C., Gaudenz, K., Peuß, R., Sung, J. Y., Martineau, B., Peavey, M., Box, A. C., Tabin, J. A., McGaugh, S., Borowsky, R., Tabin, C. J., & Rohner, N. (2018). Insulin resistance in cavefish as an adaptation to a nutrient-limited environment. Nature, 555(7698), 647–651. https://doi.org/10.1038/nature26136

Roberts, B. L., & Russell, I. J. (1972). The activity of lateral-line efferent neurones in stationary and swimming dogfish. The Journal of Experimental Biology, 57(2), 435–448.

Russell, I. J., & Roberts, B. L. (1972). Inhibition of Spontaneous Lateral-Line Activity by Efferent Nerve Stimulation. Journal of Experimental Biology, 57(1), 77–82.

RW Mitchell, Russell, W., & Elliot, W. (1997). Mexican eyeless characin fishes, genus Astyanax: Environment, distribution, and evolution. Texas Tech Press. https://scholar.google.com/scholar_lookup?title=Mexican%20eyeless%20characin%20fishes%2C%20genus%20Astyanax%3A%20environment%2C%20distribution%2C%20and%20evolution.%2C%20Special%20publications%20the%20museum%20Texas%20Tech%20University&author=R.W.%20Mitchell&publication_year=1977

Sharma, S., Coombs, S., Patton, P., & De Perera, T. B. (2009). The function of wall-following behaviors in the Mexican blind cavefish and a sighted relative, the Mexican tetra (Astyanax). Journal of Comparative Physiology A, 195(3), 225–240.

Skandalis, D. A., Lunsford, E. T., & Liao, J. C. (2021). Corollary discharge enables proprioception from lateral line sensory feedback. PLoS Biology, 19(10), e3001420. https://doi.org/10.1371/journal.pbio.3001420. https://doi.org/10.1371/journal.pbio.3001420

Soustelle, L., Aigouy, B., Asensio, M.-L., & Giangrande, A. (2008). UV laser mediated cell selective destruction by confocal microscopy. Neural Development, 3(1), 11. https://doi.org/10.1186/1749-8104-3-11

Stewart, W. J., Cardenas, G. S., & McHenry, M. J. (2013). Zebrafish larvae evade predators by sensing water flow. The Journal of Experimental Biology, 216(Pt 3), 388–398. https://doi.org/10.1242/jeb.072751

Stockdale, W. T., Lemieux, M. E., Killen, A. C., Zhao, J., Hu, Z., Riepsaame, J., Hamilton, N., Kudoh, T., Riley, P. R., van Aerle, R., Yamamoto, Y., & Mommersteeg, M. T. M. (2018). Heart Regeneration in the Mexican Cavefish. Cell Reports, 25(8), 1997–2007.e7. https://doi.org/10.1016/j.celrep.2018.10.072

Straka, H., Simmers, J., & Chagnaud, B. P. (2018). A New Perspective on Predictive Motor Signaling. Current Biology, 28(5), R232–R243. https://doi.org/10.1016/j.cub.2018.01.033

Tan, D., Patton, P., & Coombs, S. (2011). Do blind cavefish have behavioral specializations for active flow-sensing? Journal of Comparative Physiology A, 197(7), 743–754.

Teyke, T. (1985). Collision with and avoidance of obstacles by blind cave fish Anoptichthys jordani (Characidae). Journal of Comparative Physiology A, 157(6), 837–843.

Teyke, T. (1990). Morphological differences in neuromasts of the blind cave fish Astyanax hubbsi and the sighted river fish Astyanax mexicanus. Brain, Behavior and Evolution, 35(1), 23–30.

Trapani, J. G., & Nicolson, T. (2011). Mechanism of spontaneous activity in afferent neurons of the zebrafish lateral-line organ. The Journal of Neuroscience: The Official Journal of the Society for Neuroscience, 31(5), 1614–1623. https://doi.org/10.1523/JNEUROSCI.3369-10.2011

Tricas, T. C., & Highstein, S. M. (1991). Action of the octavolateralis efferent system upon the lateral line of free-swimming toadfish, Opsanus tau. Journal of Comparative Physiology A, 169(1), 25–37. https://doi.org/10.1007/BF00198170

Uysal, S. A., Erden, Z., Akbayrak, T., & Demirtürk, F. (2010). Comparison of balance and gait in visually or hearing impaired children. Perceptual and Motor Skills, 111(1), 71–80.

Van Trump, W. J., Coombs, S., Duncan, K., & McHenry, M. J. (2010). Gentamicin is ototoxic to all hair cells in the fish lateral line system. Hearing Research, 261(1–2), 42–50.

Varatharasan, N., Croll, R. P., & Franz-Odendaal, T. (2009). Taste bud development and patterning in sighted and blind morphs of Astyanax mexicanus. Developmental Dynamics: An Official Publication of the American Association of Anatomists, 238(12), 3056–3064. https://doi.org/10.1002/dvdy.22144

Von Campenhausen, C., Riess, I., & Weissert, R. (1981). Detection of stationary objects by the blind cave fish Anoptichthys jordani (Characidae). Journal of Comparative Physiology, 143(3), 369–374.

Warren, W. C., Boggs, T. E., Borowsky, R., Carlson, B. M., Ferrufino, E., Gross, J. B., Hillier, L., Hu, Z., Keene, A. C., & Kenzior, A. (2021). A chromosome-level genome of Astyanax mexicanus surface fish for comparing population-specific genetic differences contributing to trait evolution. Nature Communications, 12(1), 1–12.

Wickham, H. (2007). Reshaping Data with the **reshape** Package. Journal of Statistical Software, 21(12). https://doi.org/10.18637/jss.v021.i12

Wickham, H. (2009). The split-apply-combine strategy for data analysis.

Wickham, H., François, R., Henry, L., & Müller, K. (2019). dplyr: A Grammar of Data Manipulation. R package version 0.8. 0.1. Retrieved January, 13, 2020.

Wilke, C. O. (2019). cowplot: Streamlined plot theme and plot annotations for ‘ggplot2.’ R Package Version 0.9, 4.

Windsor, S. P., Tan, D., & Montgomery, J. C. (2008). Swimming kinematics and hydrodynamic imaging in the blind Mexican cave fish (Astyanax fasciatus). Journal of Experimental Biology, 211(18), 2950–2959.

Yamamoto, Y., Byerly, M. S., Jackman, W. R., & Jeffery, W. R. (2009). Pleiotropic functions of embryonic sonic hedgehog expression link jaw and taste bud amplification with eye loss during cavefish evolution. Developmental Biology, 330(1), 200–211.

Yoffe, M., Patel, K., Palia, E., Kolawole, S., Streets, A., Haspel, G., & Soares, D. (2020). Morphological malleability of the lateral line allows for surface fish (Astyanax mexicanus) adaptation to cave environments. Journal of Experimental Zoology Part B: Molecular and Developmental Evolution, 334(7–8), 511–517. https://doi.org/10.1002/jez.b.22953

Yoshizawa, M., Gorički, Š., Soares, D., & Jeffery, W. R. (2010). Evolution of a Behavioral Shift Mediated by Superficial Neuromasts Helps Cavefish Find Food in Darkness. Current Biology, 20(18), 1631–1636. https://doi.org/10.1016/j.cub.2010.07.017

Yoshizawa, M., Jeffery, W. R., Netten, S. M. van, & McHenry, M. J. (2014). The sensitivity of lateral line receptors and their role in the behavior of Mexican blind cavefish (Astyanax mexicanus). Journal of Experimental Biology, 217(6), 886–895. https://doi.org/10.1242/jeb.094599

